# Attenuation of influenza A virus disease severity by viral co-infection in a mouse model

**DOI:** 10.1101/326546

**Authors:** Andres J. Gonzalez, Emmanuel C. Ijezie, Onesmo B. Balemba, Tanya A. Miura

**Author notes:** Corresponding Author: Tanya A. Miura.

## Abstract

Influenza viruses and rhinoviruses are responsible for a large number of acute respiratory viral infections in human populations and are detected as co-pathogens within hosts. Clinical and epidemiological studies suggest that co-infection by rhinovirus and influenza virus may reduce disease severity and that they may also interfere with each other’s spread within a host population. To determine how co-infection by these two unrelated respiratory viruses affects pathogenesis, we established a mouse model using a minor serogroup rhinovirus (RV1B) and mouse-adapted influenza A virus (PR8). Infection of mice with RV1B two days before PR8 reduced pathogenesis of mild to moderate, but not severe PR8 infections. Disease attenuation was associated with an early inflammatory response in the lungs and enhanced clearance of PR8. However, co-infection by RV1B did not reduce PR8 viral loads early in infection or inhibit replication of PR8 within respiratory epithelia or *in vitro*. Inflammation in co-infected mice remained focal, in comparison to diffuse inflammation and damage in the lungs of mice infected by PR8. These findings suggest that RV1B stimulates an early immune response that clears PR8 while limiting excessive pulmonary inflammation. The timing of RV1B co-infection was a critical determinant of protection, suggesting that sufficient time is needed to induce this response. Finally, disease attenuation was not unique to RV1B: co-infection by a murine coronavirus two days before PR8 also reduced disease severity. This model will be critical for understanding the mechanisms responsible for attenuation of influenza disease during co-infection by unrelated respiratory viruses.

## Importance

Viral infections in the respiratory tract can cause severe disease and are responsible for a majority of pediatric hospitalizations. Molecular diagnostics have revealed that approximately 20% of these patients are infected by more than one unrelated viral pathogen. To understand how viral co-infection affects disease severity, we inoculated mice with a mild viral pathogen followed two days later by a virulent viral pathogen. This model demonstrated that rhinovirus can reduce the severity of influenza A virus, which corresponded with an early but controlled inflammatory response in the lungs and early clearance of influenza A virus. We further determined the dose and timing parameters that were important for effective disease attenuation and showed that influenza disease is also reduced by co-infection with a murine coronavirus. These findings demonstrate that co-infecting viruses can alter immune responses and pathogenesis in the respiratory tract.

## Introduction

Respiratory tract infections are a leading cause of morbidity and mortality worldwide and viruses from many different families contribute to the disease burden. Advances in viral diagnostics and surveillance have led to the finding that viral co-infection, or the presence of multiple unrelated viruses, is quite common in the respiratory tract (1–3). Viruses involved in coinfections have the potential for interactions within the host that could affect replication dynamics, immune responses, and disease pathogenesis (4, 5).

Influenza viruses and rhinoviruses are common causes of both upper and lower respiratory tract infections across all age groups (6–8). Thus, it is not surprising that they are frequently detected in co-infected patients. During the 2009 pandemic, the severity of influenza was reduced by co-infection with rhinoviruses, though it was enhanced by co-infection with other respiratory viruses (9). These differences did not correspond with changes in viral replication, as equivalent H1N1 viral loads were detected in co-infected patients and those infected by influenza A virus alone (9). Other clinical studies have found that respiratory viral co-infection can enhance (10–12), reduce (13, 14), or have no effect (15) on the severity of influenza. These differences may be due to the viruses involved, the order that they infected the host, and differences in patient populations. The disease severity of rhinovirus infections has also been shown to be affected by viral co-infections. Co-infection by influenza viruses has been found to reduce the severity of rhinovirus infection (11) and co-infection by respiratory syncytial virus enhances disease compared to rhinovirus alone (11, 16–18). Finally, epidemiological studies suggest that rhinoviruses may interfere with the spread of influenza within populations (8, 19–22). Likewise, spread of the 2009 pandemic H1N1 strain appears to have delayed the circulation of other seasonal respiratory viruses (7). Altogether, these studies suggest that respiratory viruses have complex interactions at the host and population levels. Model systems in which the viral strains and infection timing and doses are controlled are needed to understand these interactions at a mechanistic level.

In order to study the effects of viral co-infection on influenza disease pathogenesis, we established a mouse model of co-infection using mouse-adapted influenza A virus, strain PR8. PR8 infection in BALB/c mice causes dose-dependent disease severity, in which mortality corresponds with early viral replication and cytokine production, followed by massive cellular infiltration in the lungs (23–25). To determine how PR8-mediated disease is altered by a mild viral infection, we co-infected mice with human rhinovirus strain 1B (RV1B) or a murine coronavirus, mouse hepatitis virus (MHV-1). RV1B infects mice using low-density lipoprotein as its receptor, which is found on murine respiratory epithelial cells (26). Inoculation of BALB/c mice with an extremely high dose of RV1B results in neutrophil and lymphocyte recruitment to the airways, along with expression of antiviral and proinflammatory cytokines, without causing overt disease (27). MHV-1 is a respiratory tropic coronavirus that is a native pathogen of mice. While MHV-1 causes severe disease in A/J and TLR4-deficient mouse strains, it causes dose-dependent severity in the respiratory tract of BALB/c mice (28, 29).

Here, we show that RV1B (RV) reduced the pathogenesis of PR8 infection and that the severity of PR8 infection and the timing of co-infection were critical determinants of disease attenuation. Co-infection-mediated protection corresponded with early but controlled pulmonary inflammation and enhanced recovery of co-infected mice. RV1B did not prevent infection by PR8 or inhibit early replication, but led to faster clearance. Interestingly, attenuation of PR8 disease was not limited to co-infection by RV1B, it was also seen when mice were co-infected by MHV. By identifying the parameters that are critical for disease attenuation during coinfection, such as viral doses, order, and timing of infection, we have established a model system in which we can determine how co-infecting viruses interact with the host’s immune system to alter pathogenesis.

## Results

### RV infection two days before PR8 lessens disease severity in mice

Patients that are co-infected by H1N1 influenza A virus and rhinovirus have less severe disease than those that are co-infected by H1N1 and a non-rhinovirus (9). To evaluate the effects of rhinovirus co-infection on influenza disease severity, we inoculated mice with RV followed two days later by a low, medium, or high dose of PR8 (PR8_Low_, PR8_Med_, or PR8_Hi_). Control groups were inoculated with RV/mock or mock/PR8 on day −2/day 0. Mice were monitored daily for mortality and weight loss, and were given a score based on clinical signs of disease, including ruffled fur, hunched posture, lethargy, and labored breathing. Humane endpoints included loss of more than 25% of their starting weight and/or severe clinical signs of disease. Similar to previous studies (27), mice inoculated with RV had no mortality or morbidity, including weight loss and clinical signs (Fig. 1A, B, C; RV/Mock). Mice that were inoculated with RV had no distinguishable differences in weight loss or clinical signs from mice that received mock inoculations (data not shown). Mice inoculated with PR8_Low_ (Mock/PR8_Low_) reached 40% mortality by day 8, but the remaining mice survived until end of the study (Fig. 1A). These mice began losing weight and showing clinical signs on day 3, which progressed until the peak of disease on day 8 (Fig. 1B, C). Clinical signs of disease included minor ruffling of fur and hunching of the back, which progressed to include slightly labored breathing until eventual recovery from infection. In contrast to the mice infected with PR8_Low_ alone, mice inoculated with RV two days before PR8_Low_ (RV/PR8_Low_) all survived until the end of the study (Fig. 1A). Coinfected mice lost weight between days 3–l7, but the rate of loss and maximum weight loss were less than mice infected by PR8_Low_ alone (Fig. 1B). Clinical signs in co-infected mice were limited to minor ruffling and hunching and the onset of these signs was delayed until day 6 compared to mice infected with low dose PR8 alone (Fig. 1C).

**FIG 1.**
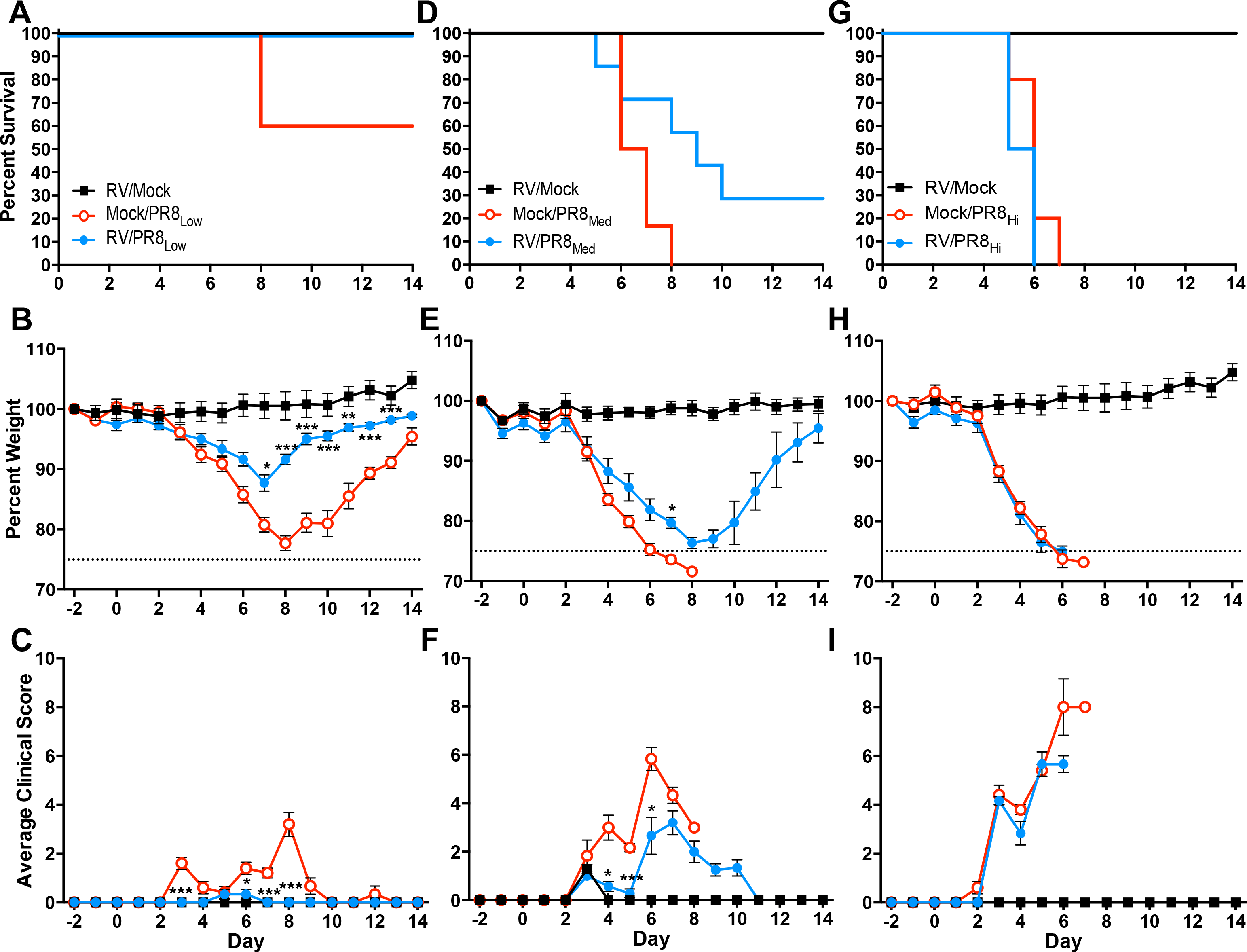
Disease kinetics in mice infected by PR8 or co-infected with RV two days before PR8. Mice were either mock-inoculated or inoculated with 7.6×10^6^ TCID_50_ units of RV on day −2. On day 0, mice were either mock-inoculated or inoculated with PR8 at (A-C) ~100 (PR8_Low_), (D-F) ~200 (PR8_Med_), or (G-I) ~7.5×10^3^ (PR8_Hi_) TCID_50_ units. Mice were monitored for (A, D, G) mortality, (B, E, H) weight loss, and (C, F, I) clinical signs of disease, including lethargy, ruffled fur, hunched back, and labored breathing. Clinical scores were assigned on a scale of 0–l3 in each category and total daily scores were averaged. Data represent the means and standard errors for 5–l7 mice and are representative of two independent experiments. Survival curves were compared using log-rank Mantel-Cox curve comparison. Weight loss and clinical score data were compared using multiple student’s *t* tests with Holm-Sidak multiple comparison correction. Significant differences compared to Mock/PR8 are indicated *P < 0.05, **P < 0.01, *** P < 0.001.

RV-mediated disease attenuation was less effective when the dose of PR8 was increased. Mice that were inoculated with a medium dose of PR8 (Mock/PR8_Med_) had increased mortality, weight loss, and clinical signs compared to Mock/PR8_Low_. Furthermore, disease severity was only partially reduced by co-infection with RV (RV/PR8_Med_). All mice inoculated with PR8_Med_ alone succumbed to infection, whereas 2 of the 7 co-infected mice survived (Fig. 1D). Both groups of mice, Mock/PR8_Med_ and RV/PR8_Med_, began losing weight on day 3 and continued to day 8 (Fig. 1E). Mice inoculated with PR8_Med_ alone had more severe clinical signs throughout PR8 infection, exhibited as severe ruffling, mild lethargy, hunching, and labored breathing (Fig. 1F). In contrast, co-infected (RV/PR8_Med_) mice had milder clinical signs limited to mild ruffling, slight hunching, and lethargy, which resolved as the surviving mice began to regain weight.

RV did not alter disease severity when mice were infected with a high dose of PR8. All of the mice inoculated with a PR8_Hi_ alone reached humane endpoints by day 7, one day earlier than mice inoculated with PR8_Med_ (Fig. 1G vs. 1D). Mock/PR8_Hi_-infected mice lost weight at a similar rate to mice to inoculated with Mock/PR8_Med_ (Fig. 1H vs. 1E) but displayed more severe clinical signs at later times in infection (Fig. 1I vs. 1F). Mock/PR8_Hi_-infected mice had moderate ruffling and hunching on day 3, which quickly progressed to severe ruffling and hunching and moderate-severe lethargy and labored breathing before succumbing to infection (Fig. 1I). Coinfected mice (RV/PR8_Hi_) had no differences in mortality (Fig. 1G), weight loss (Fig. 1H), or clinical signs (Fig. 1I) compared to mock/PR8_Hi_-infected mice. In summary, RV lessened the severity of a mild or moderate, but not severe, infection by PR8 when inoculated two days before PR8.

### Co-infection by RV leads to more rapid clearance of PR8 from the lungs

The above experiments established that RV effectively attenuated disease due to a mild PR8 infection. We next asked whether the reduced virulence during co-infection was due to inhibition of PR8 replication within the lungs. Groups of mice were either mock-inoculated or inoculated with RV two days before inoculating with a low dose of PR8 (Mock/PR8, RV/PR8). Groups of mice were euthanized on days 4, 7, and 10 after PR8 inoculation and lungs were harvested for PR8 quantification. On day 4 after PR8 inoculation, the viral loads in single virus and co-infected mice were not significantly different, but co-infected mice had greater variation within the group (Fig. 2). However, on day 7, the Mock/PR8-infected mice still had PR8 titers in the 10^4^−10^5^ range, whereas the RV/PR8-co-infected mice had undetectable levels of PR8. By day 10, both groups had undetectable levels of infectious PR8. This suggests that co-infection with RV did not prevent infection or inhibit early replication of PR8, but induced more rapid clearance of PR8 from the lungs.

**FIG 2.**
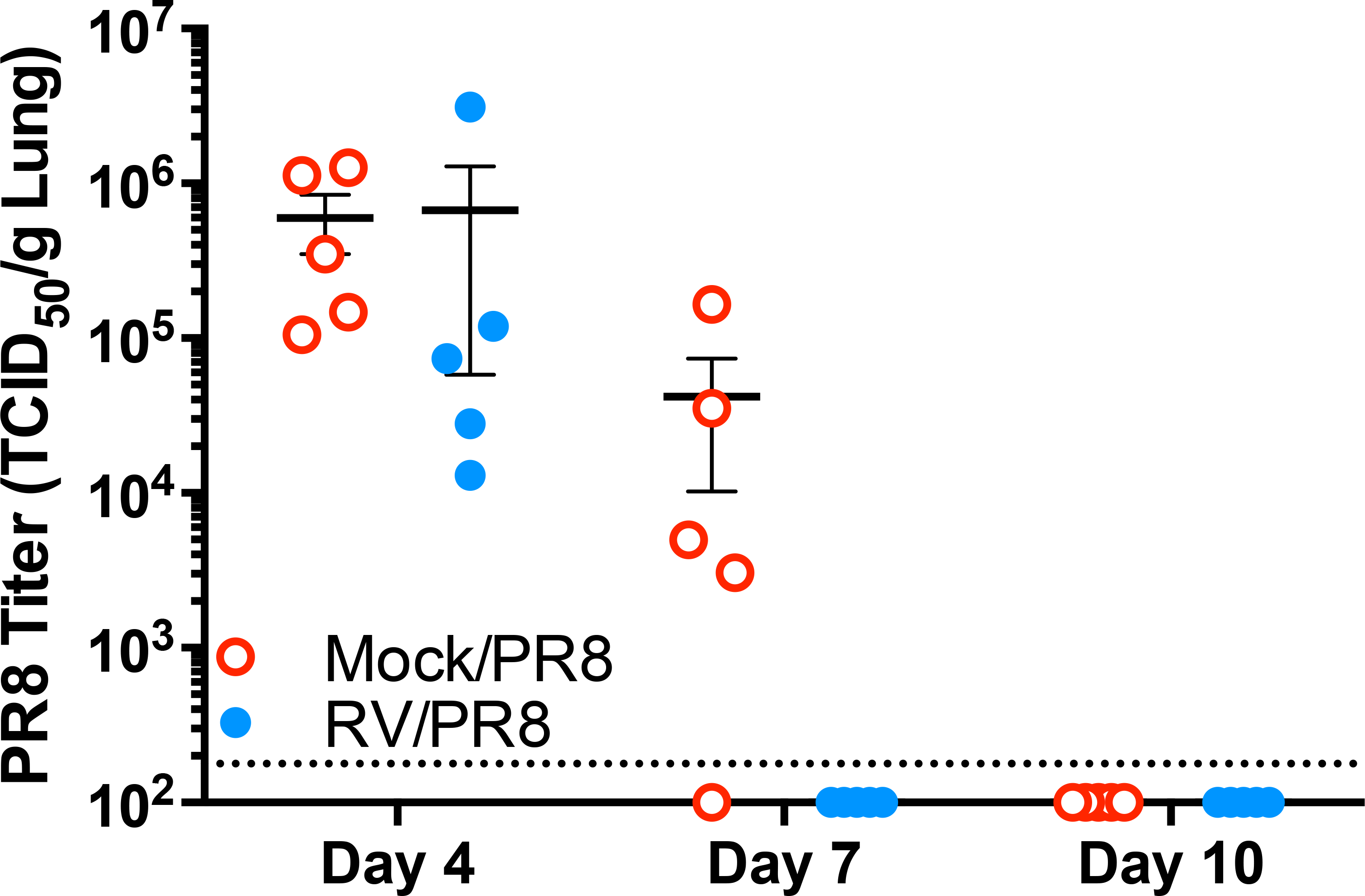
PR8 titers in the lungs of mice infected by PR8 or co-infected by RV two days before PR8. Mice were either mock-inoculated or inoculated with 7.6×10^6^ TCID_50_ units of RV on day −2. On day 0, mice were either mock-inoculated or inoculated with ~100 TCID_50_ units of PR8. PR8 was titrated by TCID_50_ assay of homogenized lungs. Data for individual mice are shown with lines indicating means and standard errors for each group. The dotted line indicates the limit of detection of the assay. Titers were compared between groups using a student’s *t*-test, which determined they were not significantly different.

We also evaluated lung sections for PR8 antigens by immunohistochemistry. The lack of PR8 antigen staining in lung sections from mock-inoculated mice confirmed the specificity of staining (Fig. 3A). On day 4, both single-and co-infected groups had extensive infection of the bronchial epithelium (not shown) in addition to viral antigen in bronchiolar epithelium and alveoli (Fig. 3B). The Mock/PR8-infected lung tissues had more extensive sloughing of infected epithelia with viral antigens associated with mucopurulent material in the airways. While Mock/PR8-infected animals had clear progression of infection from upper and lower airways into the adjacent alveoli, co-infected animals appeared to have viral antigen in the alveoli earlier and it was not always associated with infection of bronchiolar epithelium in the same region. Both groups had extensive viral antigen staining in the alveoli, with antigen detected in pneumocytes and immune cells, especially macrophages and neutrophils. On day 7, antigen staining was reduced in lung tissues from both groups and was localized in airways, especially cells that had been sloughed from the epithelium (Fig. 3C). PR8 antigens in the alveoli were mainly found in immune cells by day 7, suggesting clearance of infection. In agreement with the undetectable levels of infectious PR8, both groups had little antigen staining on day 10, which was predominantly in immune cells within focal areas (not shown). These findings support our observation that co-infection by RV did not completely inhibit replication of PR8 in the respiratory tract.

**FIG 3.**
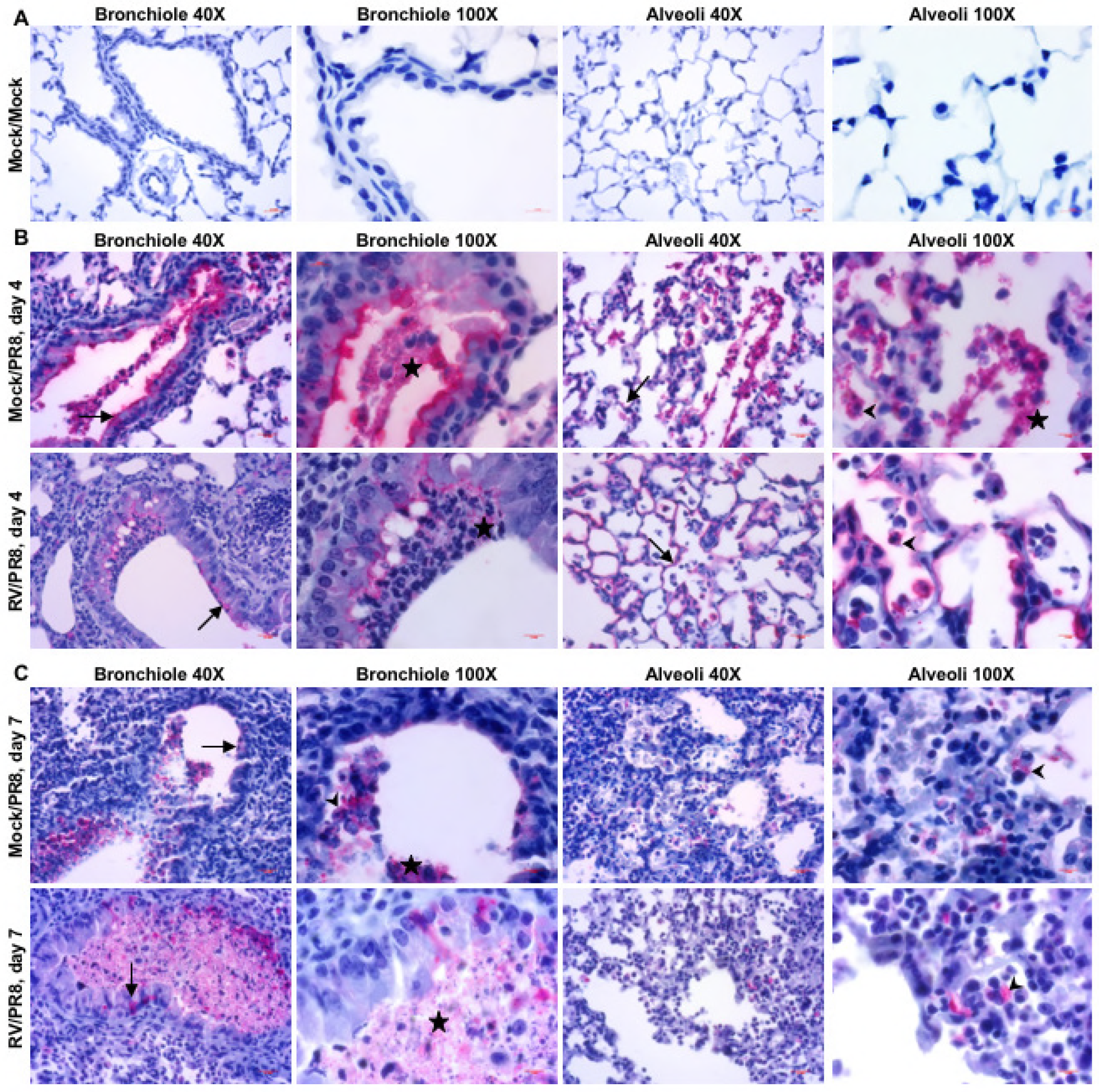
Immunohistochemistry of PR8 antigen in the lungs of mice infected by PR8 or coinfected with RV two days before PR8. Images taken in the indicated regions of the lungs and at the indicated magnifications show (A) mice given mock inoculations on days −2 and 0 and (B, C) mice given Mock/PR8 or RV/PR8 on day −2/day 0. Tissue sections were immunostained for the PR8 hemagglutinin protein, which was visualized by vector red, and counterstained with hematoxylin. Lung tissues were collected on (B) day 4 and (C) day 7 after PR8 inoculation. Images were representative of multiple sections from two animals per group and time point. Arrows show examples of antigen in epithelial cells; arrowheads show examples of antigen in leukocytes, predominantly macrophages and neutrophils; stars indicate mucopurulent material.

### RV does not interfere with replication of PR8 *in vitro*

Next, we used a murine lung epithelial cell line (LA-4) that is susceptible to infection by both viruses (30) to determine if co-infection by RV would interfere with PR8 replication *in vitro*. LA-4 cells were inoculated with RV 6 or 12 hours before or simultaneously with PR8. PR8 released into the media was quantified by TCID_50_ assay in MDCK cells. We confirmed that the presence of RV in these samples did not interfere with quantification of PR8 in MDCK cells (not shown). There were no significant differences at any time point between cells inoculated with PR8 alone or co-infected with RV either simultaneously with or 12 hours before PR8 (Fig. 4A, C). There were significant differences at 24 and 96 hours when cells were inoculated with RV 6 hours before PR8 (Fig. 4B). The lower PR8 titers at 96 hours may have been due to higher cell death from two viruses being present, providing fewer susceptible cells for PR8 replication. Though not significant, this trend was also seen when cells were inoculated with both viruses simultaneously (Fig. 4A). These data help corroborate our *in vivo* finding that RV did not inhibit replication of PR8. Rather, co-infection is most likely stimulating the immune system, leading to faster clearance of PR8.

**FIG 4.**
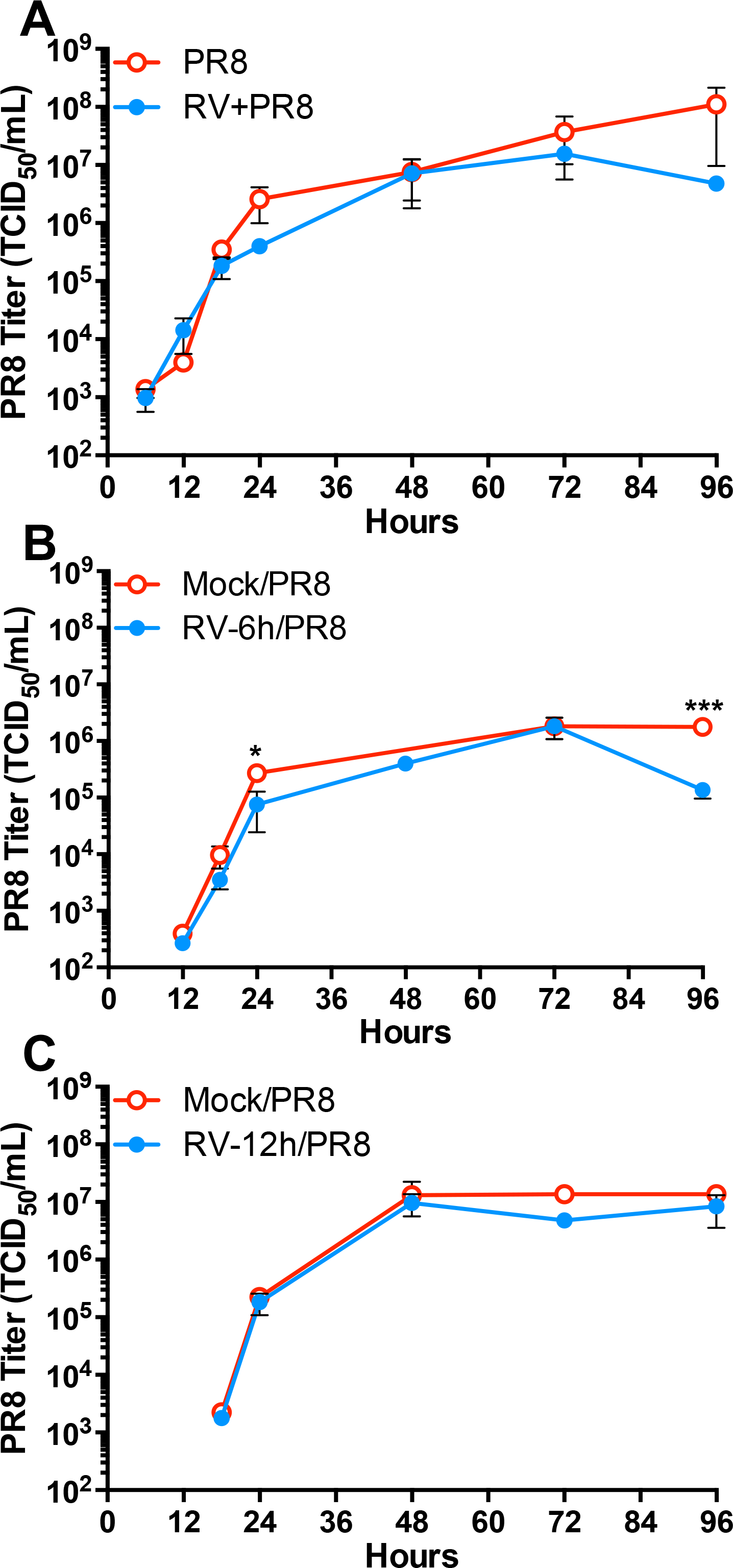
Growth curves of PR8 from cells infected by PR8 or RV and PR8. LA-4 cells were inoculated with RV (moi = 1) (A) simultaneously with, (B) 6 hours before, or (C) 12 hours before inoculation with PR8 (moi =1). Media was collected at the indicated times after PR8 inoculation and titrated for PR8 by TCID_50_ assay. Significant differences compared to Mock/PR8 were determined by students *t*-test. *P < 0.05, **P < 0.01, *** P < 0.001.

### Co-infection with RV stimulated an early yet controlled inflammatory response to PR8 infection

We next evaluated the histopathology of lungs from single-(Mock/PR8) and co-(RV/PR8) infected mice on days 4, 7, and 10 after inoculation with PR8. Compared to mock-inoculated controls, both Mock/PR8-and RV/PR8-infected mice had multifocal tracheobronchitis and bronchioalveolitis (Fig. 5 A-C and not shown). Lung pathology was characterized by epithelial degeneration, necrosis and sloughing, and accumulation of neutrophils, macrophages, and lymphocytes in the inflamed areas. This was associated with the accumulation of mucopurulent discharge in the bronchioles and alveoli (Fig. 5C). The extension of inflammation from bronchioles into the surrounding lung parenchyma resulted in localized alveolitis. In general, inflammation was associated with dilated and congested blood capillaries with extravasation of blood plasma, hemorrhage, thickened alveolar septae, collapsed alveoli, and enlarged alveolar ducts observed in severe lesions (Fig. 5B, C). Although focal inflammation was found in all lobes, the right lung’s caval lobe appeared to be commonly more inflamed than other lobes. Pulmonary invagination was often associated with inflamed portions of the lungs.

**FIG 5.**
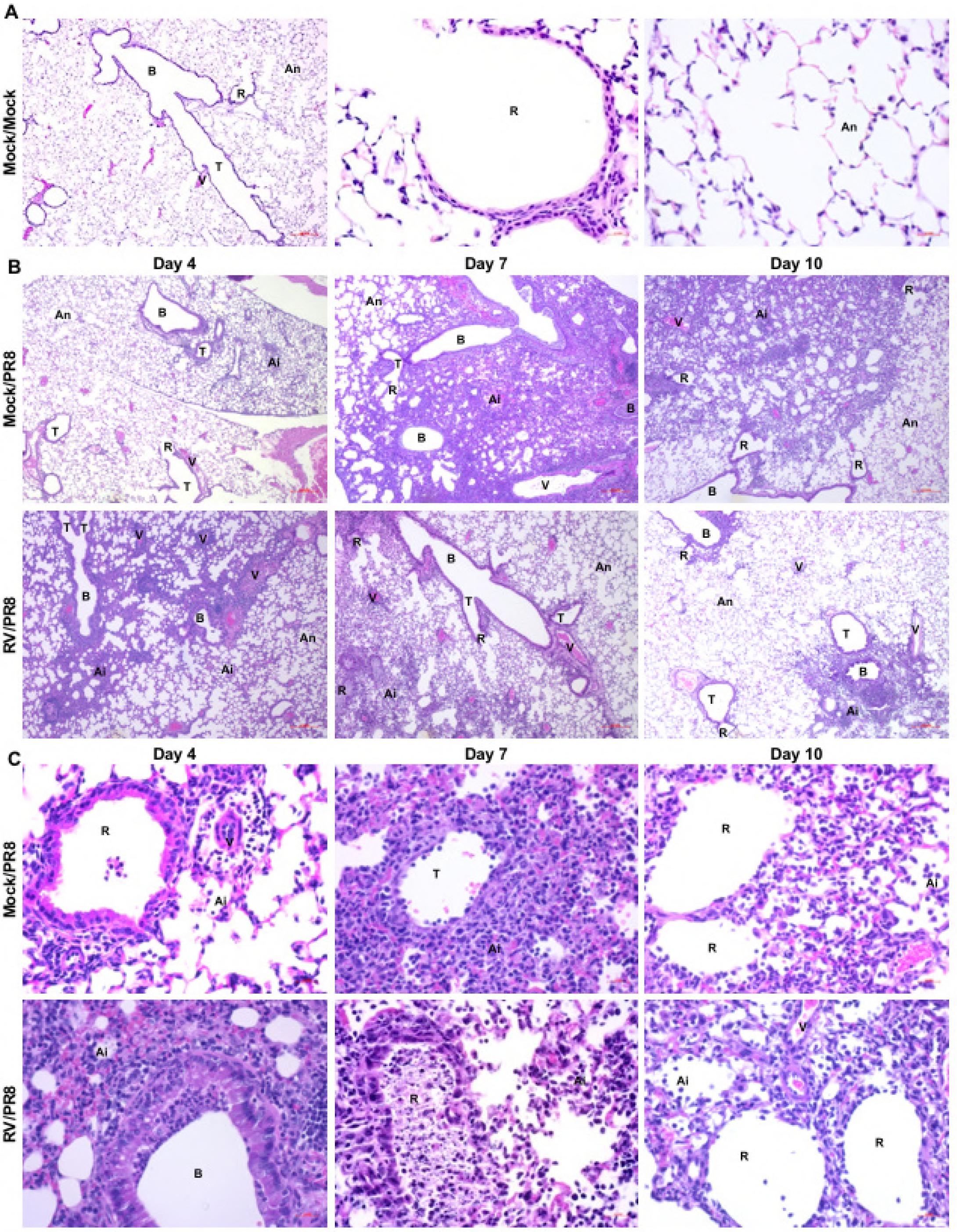
Histopathology of mouse lungs infected by PR8 or co-infected with RV two days before PR8. Mice were either mock-inoculated or inoculated with 7.6×10^6^ TCID_50_ units of RV on day - 2. On day 0, mice were either mock-inoculated or inoculated with ~100 TCID_50_ units of PR8. Lungs were paraffin-embedded and sections were stained with hematoxylin and eosin. Images were representative of multiple tissue sections from two mice per group and time point. (A) shows images from lung sections of Mock/Mock inoculated mice taken with 10X and 40X objectives. (B) shows images from the indicated groups and days taken with 10X objective. (C) shows images from the indicated groups and days taken with 40X objective. Labeled structures include bronchioles (B), terminal bronchioles (T), respiratory bronchioles (R), normal alveoli (An), inflamed alveoli (Ai), and blood vessels (V).

Co-infection with RV induced earlier inflammation, but reduced the severity of inflammation elicited by PR8. By day 4, lung pathology was more extensive in RV/PR8 coinfected mice than Mock/PR8-infected mice. However, by days 7 and 10, mice infected with Mock/PR8 had more severe lung pathology than RV/PR8 co-infected mice. Overall, necrosis and desquamation in the trachea, bronchi, bronchioles, and alveoli were more severe in Mock/PR8 than in RV/PR8 co-infected mice. Furthermore, excessive mucopurulent material consisting of neutrophils, macrophages, cellular debris, and transudate accumulated in the lungs of Mock/PR8-infected mice. Congestion and hemorrhage were also more severe and pleurisy was only present in the lungs of Mock/PR8-infected mice. Furthermore, the expansion of bronchiolar inflammation into the parenchyma and collapse and destruction of alveoli was less extensive in lungs of RV/PR8 co-infected mice. Noteworthy, perivascular cuffing was more dense and widespread in RV/PR8 co-infected mice and their lungs had more lymphocytes and macrophages and fewer neutrophils than Mock/PR8-infected mice. These findings suggest that Mock/PR8-infected mice had more tissue damage and exacerbated inflammation while RV/PR8 co-infected mice had a more focused anti-viral cellular immune response.

Co-infection by RV also resulted in earlier resolution of inflammation and tissue regeneration. Type II pneumocyte hyperplasia and regeneration of the epithelium was observed in the bronchioles and alveoli of RV/PR8 co-infected mice on days 7 and 10, indicating signs of regeneration. Overall, there were fewer inflammatory cells in the alveoli in lungs of RV/PR8 coinfected mice compared with lungs of Mock/PR8-infected mice. Pleurisy was encountered in some of the lungs obtained from Mock/PR8-infected mice but not RV/PR8 co-infected mice. In summary, the findings of this study show that co-infecting mice with RV reduced the magnitude of the inflammatory response to PR8 infection and accelerated epithelial repair and alveoli restoration, and the overall recovery.

### The timing of RV co-infection determines the effect on disease severity during PR8 infection

Co-infection by RV was effective at reducing disease during a mild or moderate PR8 infection when given two days before PR8. We next determined whether co-infection with RV simultaneously with or two days after a low or medium dose of PR8 would also ameliorate disease. When RV and PR8_Low_ were inoculated simultaneously (RV+PR8_Low_), co-infection resulted in a disease phenotype intermediate between PR8_Low_ alone (Mock/PR8_Low_) and RV two days before PR8_Low_ (RV/PR8_Low_) (Fig. 6A-C). RV+PR8_Low_ co-infected mice reached 33% mortality by day 9 (Fig. 6A). This mortality was higher than mice inoculated with RV two days before PR8_Low_ (RV/PR8_Low_), but significantly lower than the 100% mortality seen with mice infected with PR8_Low_ alone (Mock/PR8_Low_). RV+PR8_Low_-infected mice lost weight at a similar rate to Mock/PR8_Low_-infected mice, beginning on day 3 and continuing until day 8 before recovering (Fig. 6B). Average clinical scores were also similar between simultaneous co-infection and single infection, with slight lethargy, hunching, breathing, and moderate ruffling detected on day 3 and 4 and peaking with moderate lethargy, hunching, breathing, and severe ruffling on day 7 before co-infected mice (RV+PR8_Low_) began recovering from infection (Fig. 6C). Interestingly, when RV was given two days after PR8_Low_ (PR8_Low_/RV), co-infection exacerbated PR8 disease, as evidenced by more rapid mortality, weight loss, and higher clinical scores than mice infected with Mock/PR8_Low_ (Fig. 6A-C). PR8_Low_/RV co-infected mice began dying and reached 100% mortality two days before mice inoculated with Mock/PR8_Low_ (Fig. 6B).

**FIG 6.**
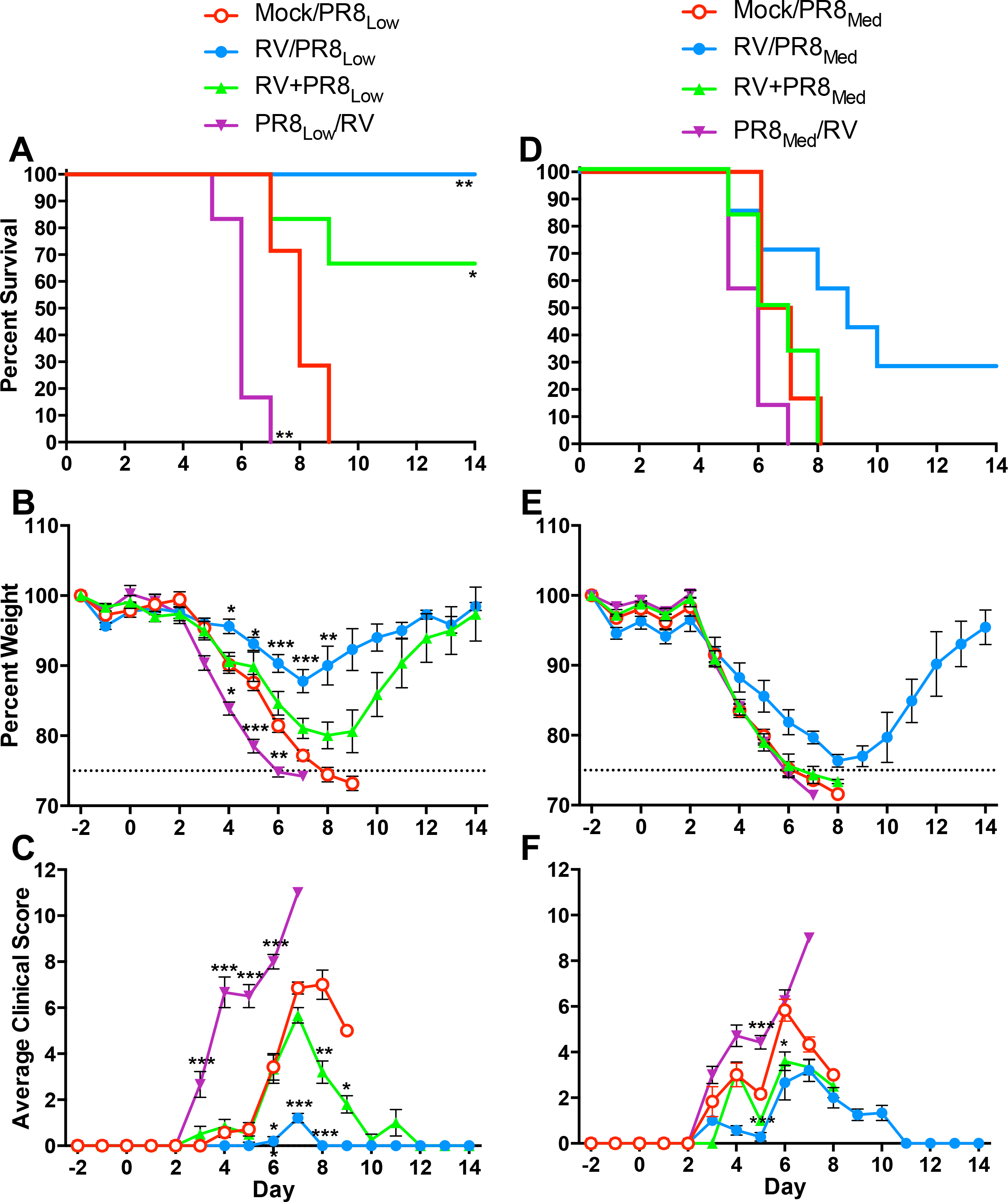
Disease kinetics in mice co-infected by RV two days before, simultaneously with, or two days after PR8. Groups of 6–l7 BALB/c mice were either mock-inoculated or inoculated with 7.6×10^6^ TCID_50_ units of RV two days before (RV/PR8), simultaneously with (RV+PR8), or two days after (PR8/RV) either inoculation with ~100 (PR8_Low_) or ~200 (PR8_Med_) TCID_50_ units of PR8. Mice were monitored for mortality, weight loss, and clinical signs of disease (lethargy, ruffled fur, hunched posture, labored breathing) for 14 days after PR8 inoculation. (A-C) RV and low dose PR8 co-infection mortality (A), weight loss (B), and clinical scores (C). (D-F) RV and medium dose PR8 co-infection mortality (D), weight loss (E), and clinical scores (F). Survival curves were compared using log-rank Mantel-Cox curve comparison. Weight loss and clinical score data were compared using multiple student’s *t* tests with Holm-Sidak multiple comparison correction. Significant differences compared to Mock/PR8 are indicated *P < 0.05, **P < 0.01, *** P < 0.001.

PR8_Low_/RV co-infected mice also lost weight at a greater rate and had dramatically higher clinical scores than other single virus-and co-infected groups (Fig. 6B, C). These increased clinical scores were due to severe lethargy and ruffling, moderate hunching, and labored breathing, which occurred earlier during infection compared to mice in the other groups (Fig. 6C).

In contrast to PR8_Low_, RV was only effective at disease attenuation when given two days before a higher dose (PR8_Med_) of PR8. There were no significant differences in mortality, weight loss, or clinical signs between simultaneously co-infected (RV+PR8_Med_) mice and mice inoculated with PR8_Med_ alone (Fig. 6D-F). Both of these groups steadily lost weight between days 3–l8 and reached 100% mortality by day 8 (Fig. 6D, E). RV+PR8_Med_ co-infected mice had slightly lower clinical scores than Mock/PR8_Med_-infected mice (Fig. 6F), and did not experience severely labored breathing like Mock/PR8_Med_-infected mice. However, these differences in clinical signs did not affect mortality. When RV was inoculated two days after PR8_Med_ (PR8_Med_/RV), all mice reached humane endpoints by day 7, one day earlier than mice inoculated with Mock/PR8_Med_ (Fig. 6D). PR8_Med_/RV co-infected mice lost weight at a similar rate to Mock/PR8_Med_-infected mice and had similar clinical scores (Fig.6E-F). These data demonstrate that RV provides protection from PR8-induced disease in a time-dependent manner. Disease attenuation was most effective when RV had adequate time to activate a protective response.

### Disease attenuation is not limited to co-infection by RV

We next evaluated whether attenuation of PR8 disease was limited to co-infection by RV or if a different respiratory virus would also be effective. Mice were either mock-inoculated or inoculated with MHV two days before PR8 and monitored for survival and weight loss over 14 days. One mouse inoculated with 2×10^3^ PFU of MHV (MHV_2000_/Mock) reached a humane endpoint, but the remaining mice survived until the end of the study (Fig. 7A). MHV-infected mice (MHV_2000_/Mock) had rapid, early weight loss followed by gradual recovery of weight (Fig. 7B). These mice had moderate ruffling and hunching early in infection, which reduced to slight ruffling, lethargy, and hunching before finally recovering on day 13 (Fig. 7C). Despite the prolonged morbidity in MHV-infected mice, mice co-infected with a lethal dose of PR8 (MHV_2000_/PR8) had similar mortality and morbidity as those infected by MHV alone (Fig. 7A-C). MHV_2000_/PR8 co-infected mice regained weight at a slower rate than mice inoculated with MHV_2000_/Mock, but eventually reached the same weight (Fig. 7B). Thus, co-infection by MHV reduced the severity of PR8-induced disease, but this dose of MHV caused significant morbidity in BALB/c mice.

**FIG 7.**
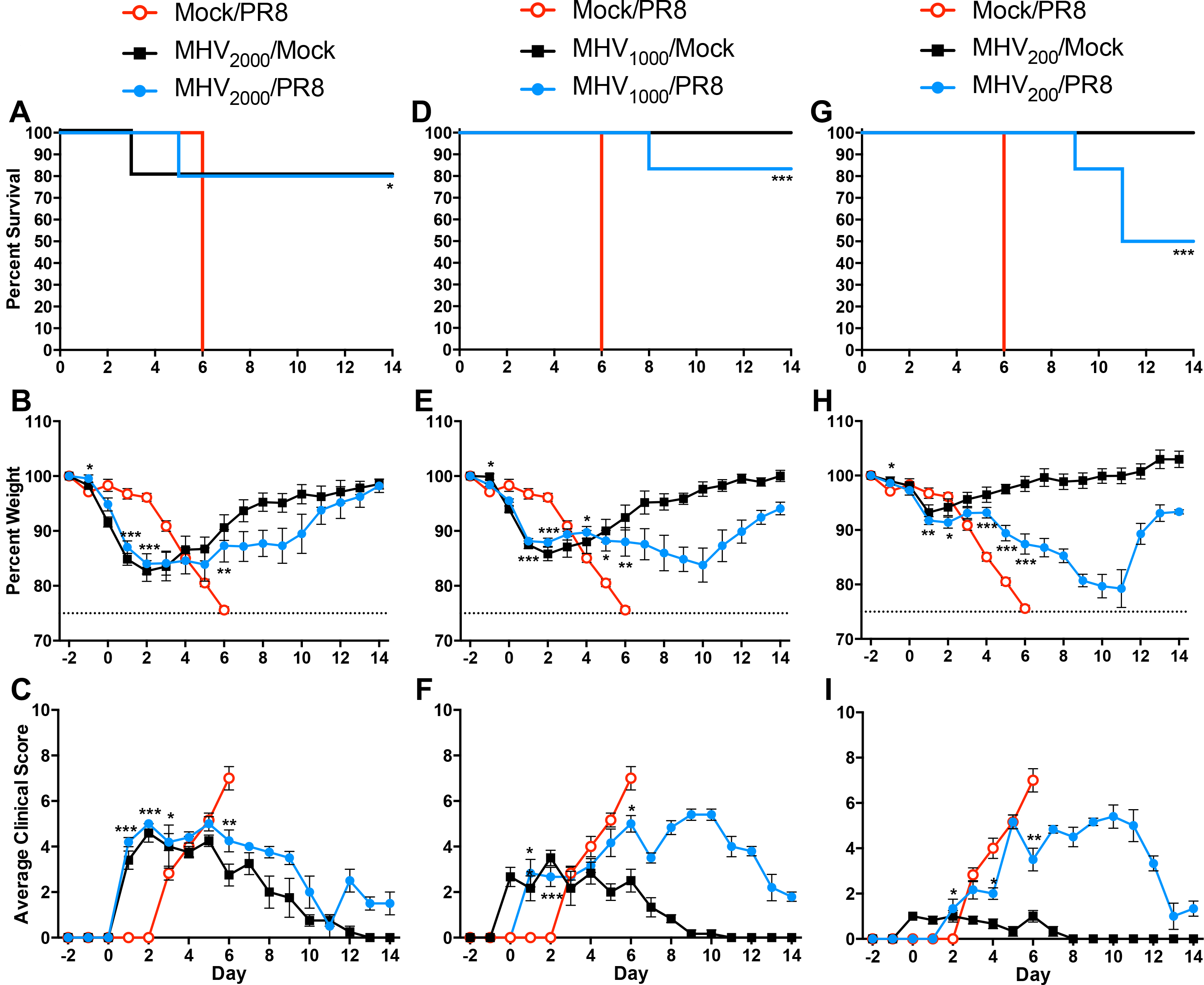
Disease kinetics in mice co-infected by MHV two days before PR8. Groups of 5–l6 BALB/c mice were either mock-inoculated or inoculated with 2.0×10^3^ (MHV_2000_), 1.0×10^3^ (MHV_1000_), or 2.0×10^2^ (MHV_200_) PFU of MHV two days before inoculation with ~100 TCID_50_ units of PR8. Mice were monitored for mortality, weight loss, and clinical signs of disease (lethargy, ruffled fur, hunched posture, labored breathing) for 14 days after PR8 inoculation. (AC) MHV_2000_ and PR8 co-infection mortality (A), weight loss (B), and clinical scores (C). (D-F) MHV_1000_ and PR8 co-infection mortality (D), weight loss (E), and clinical scores (F). (G-I) MHV_200_ and PR8 co-infection mortality (G), weight loss (H), and clinical scores (I). Survival curves were compared using log-rank Mantel-Cox curve comparison. Weight loss and clinical score data were compared using multiple student’s *t* tests with Holm-Sidak multiple comparison correction. Significant differences compared to Mock/PR8 are indicated *P < 0.05, **P < 0.01, *** P < 0.001.

To determine whether infection by a lower dose of MHV would cause less disease but still provide protection against PR8, we tested two additional doses of MHV: 1×10^3^ PFU (MHV_1000_) and 2×10^2^ PFU (MHV_200_). When mice were inoculated with the 2-fold lower dose of MHV (MHV_1000_/Mock), there was no mortality associated with infection (Fig. 7D). These mice lost weight very early in infection, but did not lose as much weight as those infected by the higher dose (Fig. 7E vs. B). MHV_1000_/Mock-infected mice had minor ruffling and hunching on day 0, which continued until day 9 (Fig. 7F). When MHV was inoculated two days before PR8 (MHV_1000_/PR8), only one mouse reached a humane endpoint, and mortality was delayed until day 8, compared to day 5 for the higher dose of MHV (Fig. 7D vs. A). Similar to mice infected by MHV_1000_ alone, MHV_1000_/PR8 co-infected mice lost weight early. However, co-infected mice had delayed recovery of weight and their weight was still lower than MHV_1000_-infected mice at the end of the study (Fig. 7E), which corresponded with prolonged clinical signs of disease (Fig. 7F). Although this lower dose of MHV caused a milder infection and provided similar protection against PR8-mediated mortality, morbidity was prolonged in the co-infected mice suggesting it was less effective than the higher dose.

When mice were inoculated with a 10-fold lower dose of MHV (MHV_200_/Mock), they all survived and had minimal weight loss and clinical signs of disease, which was limited to slight ruffling (Fig. 7G-I). However, mice that were co-infected with this low dose of MHV prior to PR8 (MHV_200_/PR8) had significant mortality, weight loss, and prolonged clinical signs of disease, including moderate lethargy and labored breathing (Fig. 6G-I). Even though this lowest dose of MHV caused minimal disease in BALB/c mice, it did not protect mice against PR8-induced morbidity and mortality. These data suggest that a dose of MHV that causes significant morbidity is required to effectively protect against a lethal dose of PR8. Although both RV and MHV given two days earlier both reduced the severity of PR8, they vary in pathogenesis and effectiveness of disease attenuation.

## Discussion

Clinical studies have suggested that respiratory viral co-infections may alter pathogenesis compared to infection by the viruses individually. RV has been implicated in both interfering with the spread of influenza viruses and reducing the severity of influenza during co-infection (8, 9, 19–22). Here, we developed a murine model of respiratory viral co-infection to study the effects on disease severity in a system where we can control the virus strains, doses, order, and timing of which they infect the host. We found that mice given RV two days before PR8 were completely protected against mortality and had reduced morbidity. RV was less effective at disease attenuation when mice were given higher doses of PR8, or RV was given at the same time as PR8. Further, mice given RV two days after PR8 had enhanced disease compared to PR8 alone. We also found that disease attenuation was not limited to co-infection by RV. A respiratory tropic strain of MHV also reduced PR8 disease when given two days before PR8. Unlike RV, MHV caused significant morbidity in mice. However, reducing the dose of MHV to lessen pathogenesis was less effective at reducing the severity of PR8. These data suggest that changes in pathogenesis during co-infection are dependent upon the severity of each infection and the order and timing of inoculation.

Despite preventing mortality and significantly inhibiting morbidity, infection of mice with RV two days before PR8 did not reduce PR8 levels in the lungs early in infection (Fig. 2) or prevent spread of PR8 within the respiratory tract (Fig. 3). These findings suggest that RV does not directly inhibit infection by PR8, which was also confirmed by our *in vitro* studies (Fig. 4). Our previous studies have shown that RV induces a robust type I IFN response in the LA-4 cell line (30), thus the lack of PR8 inhibition we saw is not due to the absence of an IFN response. This is not surprising, as the NS1 protein of PR8 is known to antagonize type I IFN responses (31). Others have shown that RV induces expression of type I and type III IFNs in the respiratory tract of infected mice (27, 32, 33). Although our data do not support a role for RV-induced IFN responses in preventing infection by PR8, IFNs may be important for inducing downstream antiviral responses that contribute to earlier clearance of PR8 in co-infected animals. In addition to promoting cell-intrinsic antiviral defense strategies, type I IFNs are important for the recruitment and functional phenotypes of myeloid cell responses to influenza virus infections (34, 35). In the absence of type I IFN signaling, PR8 disease severity is increased, but the enhanced disease is not completely due to increased viral loads. Rather, these studies showed that type I IFN signaling is needed to down-regulate inflammatory monocyte and neutrophil responses during PR8 infection. Furthermore, in the absence of type I IFN signaling, monocytes develop increased inflammatory phenotypes and reduced expression of genes that down-regulate inflammation (34, 35). Thus, type I IFN responses induced by RV could promote down-regulation of inflammation that we observed on days 7 and 10.

Histopathology analyses demonstrated that mice inoculated with RV two days before PR8 had earlier recruitment of immune cells into the lungs (Fig. 5). On day 4 after PR8 infection, co-infected animals had multiple foci of inflammation throughout the lungs. In contrast, animals infected with PR8 alone had reduced recruitment of inflammatory cells. This early immune response may lead to earlier clearance of PR8 from co-infected lungs, as we saw by day 7 (Fig. 2). Although it is not virulent in mice, RV (strain 1B) induces an inflammatory response in the respiratory tracts of BALB/c and C57Bl/6 mice. The response to RV includes recruitment of neutrophils and lymphocytes, concurrent with production of antiviral cytokines and chemokines. Neutrophil levels in the airways of RV-infected mice peak on days 1 and 2 and decline by day 4, while lymphocytes are present through day 7 (27, 32, 33, 36, 37). Studies differ in whether macrophage numbers change significantly upon RV infection in mice (27, 32, 33, 36). Type I and III interferons, proinflammatory cytokines, and neutrophil and lymphocyte recruiting chemokines are also up-regulated in response to RV infection in mice (27, 32, 33). These cellular and cytokine responses are not stimulated by UV-inactivated virus, suggesting that viral replication is required (27, 33). We observed increased macrophages and lymphocytes and enhanced perivascular cuffing in histology sections from RV/PR8 co-infected animals, suggesting a robust cellular immune response. Ongoing studies in our lab will determine the RV-induced immune components that are required for attenuation of PR8 disease during co-infection.

In contrast to the enhanced inflammation in the lungs of co-infected mice on day 4, the histopathology on days 7 and 10 was less severe in co-infected compared to PR8-infected mice (Fig. 5). This could be an indirect consequence of early viral clearance (Fig. 2) or active down-regulation of the inflammatory response in co-infected animals. Multiple studies have shown that inflammation during influenza and other respiratory viral infections can cause pathogenesis that is independent of viral levels in the lungs (34, 35, 38–40). Our histopathology data are in agreement with studies that demonstrate excessive inflammatory responses occurring as viral loads are decreasing (23, 41). We inoculated mice with PR8 two days after RV inoculation, which is just prior to the decline in neutrophil numbers in the lungs of RV-infected BALB/c mice (27). Furthermore, RV infection actively down-regulates neutrophil responses by inhibiting TLR signaling and thereby reducing expression of neutrophil-specific chemokines upon co-infection by a bacterial pathogen (42). The inflamed areas of the RV/PR8 co-infected lungs contained fewer neutrophils than mock/PR8-infected lungs. Neutrophils are known to contribute to immune-mediated damage during PR8 infection (43, 44). Thus, active down-regulation of neutrophilic inflammation by RV may contribute to the reduced severity of PR8 in our studies.

This study demonstrated that RV-mediated disease attenuation was less effective as we increased the dose of PR8, shortened the timing between virus inoculations, or gave RV after PR8 (Figs. 1 and 6). These differences are likely due to the kinetics and magnitudes of RV-and PR8-induced immune responses. Higher doses of PR8 likely overwhelm defense mechanisms that are induced by RV and these defense mechanisms are likely not induced quickly enough to completely protect against PR8 given at the same time. However, the observation that RV given two days after PR8 exacerbates pathogenesis suggests that once a response to PR8 has been initiated, the RV-induced response may aggravate immunopathology. RV induces multiple immune components that are known to contribute to influenza disease, including neutrophils (43), NK cells (45), and type I IFNs (46, 47). Further studies are needed to identify the mechanisms that exacerbate disease when RV is given 2 days after PR8. It is possible that these are the same mechanisms that reduce disease when RV is given 2 days before PR8 and that the timing of their induction is key to regulating pathogenesis.

Interestingly, infection by MHV two days prior to PR8 also reduced the severity of PR8 infection. Unlike RV, MHV causes morbidity and mortality in mice, though virulence is mouse strain-dependent (28, 29, 48). We have observed dose-dependent severity of MHV in BALB/c mice. Intranasal inoculation of 2 × 10^3^ PFU resulted in clinical disease with low mortality (Fig. 7), while a dose of 2 × 10^5^ caused 100% mortality (data not shown). Survival of BALB/c mice upon MHV-1 infection is dependent upon a type I IFN response (49). Furthermore, mouse strain-dependent resistance to MHV-1 disease corresponds with robust expression of type I IFNs (28). Based on these studies, we expect that the type I IFN response is adequate to protect BALB/c from a lower dose, but not a lethal dose, of MHV. Other components of the BALB/c immune response to MHV-1 infection have not been studied in detail. A complete understanding of this response will be important in determining the similarities and differences in the mechanisms whereby MHV and RV reduce the severity of PR8.

Although our viral inoculations were all within two days of each other, other studies have shown that infection of mice by one virus can alter the immune response to a heterologous virus given after resolution of the initial infection (50). Mice given lymphocytic choriomeningitis virus (LCMV), murine cytomegalovirus (MCMV), or PR8 six weeks prior to vaccinia virus had reduced titers of vaccinia virus compared to control mice. However, infection with PR8 six weeks prior to LCMV or MCMV increased titers of the second virus. Protection was associated with changing from a response dominated by neutrophils to lymphocytes or a Th2 response to a Th1 response. In contrast, when pigs were infected with porcine reproductive and respiratory syndrome virus they had increased severity of porcine respiratory coronavirus infection, which corresponded with an increased Th1 cytokine response and reduced NK cell and type I IFN responses (51). Characterizing the immune responses across our various co-infection pairs and infection timings will provide insight into how different viruses mediate heterologous immunity and if these mechanisms are generalizable.

## Materials and Methods

### Virus stocks and cell lines

Madin-Darby canine kidney cells (MDCK; ATCC #CCL-34), murine fibroblast line 17Cl.1 (52) (provided by Dr. Kathryn Holmes, University of Colorado Denver School of Medicine), and HeLa cells (ATCC #CCL-2) were grown in Dulbecco’s modified Eagle medium (DMEM) supplemented with 10% fetal bovine serum (FBS; Atlanta Biologicals), and 1X antibiotic-antimycotic (ThermoFisher). Murine lung epithelial cells (LA-4; ATCC #CL-196) were grown in Ham’s F12 (Kaign’s modified) media (F12K; Caisson) supplemented with 10–l15% FBS and antibiotics. PR8 (A/Puerto Rico/8/1934 (H1N1)), obtained from BEI Resources (NR-3169), was grown and titrated by tissue culture infectious dose-50% (TCID_50_) assay in MDCK cells. MHV-1 (ATCC #VR-261) was grown and titrated by plaque assay in 17Cl.1 cells. RV1B (ATCC #VR-1645) was grown and titrated by TCID_50_ assay in HeLa cells. RV1B stocks were concentrated by centrifugation through 30% sucrose and the virus pellet was resuspended in PBS/2% FBS.

### Mouse infections

All experimental protocols were approved by the University of Idaho Institutional Animal Care and Use Committee, following the National Institutes of Health Guide for the Care and Use of Laboratory Animals. As described below, mice were monitored daily and were euthanized by an overdose of sodium pentobarbital if they reached humane endpoints.

Six to eight-week-old female BALB/c mice were purchased from Harlan Laboratories/Envigo. Mice were housed in individually vented cages with controlled light/dark cycles and regulated temperature maintained by University of Idaho Lab Animal Research Facilities and received food and water *ad libitum*. Mice were acclimatized to the facility for 5–l12 days before experiments were performed under ABSL2 conditions. Mice were anesthetized with inhaled isoflurane and inoculated intranasally with 50 uL of virus. For co-infections, mice were inoculated with each virus 2 days apart or simultaneously in a total 50 uL volume. Control mice received mock inoculations of the same buffer as the respective virus: RV (PBS/2% FBS), PR8 (MEM/2% FBS) or MHV (DMEM/10%FBS). See the results sections and figure legends for viral doses used in each experiment.

Mice were weighed and observed for clinical signs of disease daily and were humanely euthanized if they lost more than 25% of their starting weight or exhibited severe clinical signs of disease. Mice were given a daily severity score of 0–l3 in each of four categories: ruffled fur, lethargy, labored breathing, and hunched posture. The daily scores were totaled for each individual mice and averaged across the group of mice.

### PR8 quantification

Right lobes of the lungs were flash-frozen and stored at −80°C. Frozen tissues were weighed and homogenized in DMEM with 2% BSA and 1% antibiotics, and PR8 was quantified by TCID_50_ assay on MDCK cells (53). RV does not interfere with titration of PR8 in co-infected samples when using the MDCK cell line (data not shown).

### Histology and immunohistochemistry

The tracheas of euthanized mice were cannulated, and the lungs were inflated with 10% formalin before submerging in 10% formalin. After fixation, lungs were embedded in paraffin, cut in 5 um sections, and stained with modified Harris hematoxylin and eosin (VWR Scientific). Lungs were processed for immunohistochemistry in the same manner as for histology. Tissue sections were deparaffinized, rehydrated, and subjected to heat-induced antigen retrieval in 10 mM sodium citrate buffer (pH 6) with 0.01% Tween 20. Endogenous peroxidase and alkaline phosphatase activities were inhibited with BLOXALL solution (Vector Laboratories). Lung sections were immunostained for the PR8 hemagglutinin protein (HA) using a polyclonal goat antibody (NR-3148, BEI Resources) and an alkaline phosphatase conjugated anti-goat antibody (ImmPRESS, Vector Laboratories) with detection by ImmPACT Vector Red (Vector Laboratories) and counterstaining with hematoxylin. Images were acquired with a Zeiss Axio Lab.A1 microscope with an Axiocam 105 color camera using ZEN 2.3 software (Zeiss).

### *In vitro* co-infection experiments

LA-4 cells were inoculated with PR8 (MOI=1) either 6 or 12 h after or simultaneously with inoculation with RV (MOI=1). After 1 h incubation with the inoculum, cells were washed twice with serum-free medium, then incubated in Ham’s F12K medium with 2% FBS and antibiotics at 37°C. Supernatant media was collected from the cells at 6, 12, 18, 24, 48, 72, and 96 hours after PR8 inoculation and PR8 was titrated by TCID_50_ assay using MDCK cells.

### Statistics

Statistical analyses were performed using Graphpad Prism 6 software. Survival curves were compared using log-rank Mantel-Cox curve comparison. Weight loss and clinical score data were compared using multiple student’s *t* tests with Holm-Sidak multiple comparison correction. PR8 titers from mouse lungs and cell culture experiments were compared using student’s *t* tests without correction for multiple comparisons.

## Acknowledgements

Research reported in this publication was supported by the National Institute of General Medical Sciences of the National Institutes of Health under Award Number P20GM104420. The content is solely the responsibility of the authors and does not necessarily represent the official views of the National Institutes of Health. The following reagents were obtained through the NIH Biodefense and Emerging Infections Research Resources Repository, NIAID, NIH: Influenza A Virus, A/Puerto Rico/8/34 (H1N1) and Polyclonal Anti-Influenza Virus H1 (H0) Hemagglutinin (HA), A/Puerto Rico/8/34 (H1N1) Antiserum, Goat (NR-3148). The authors would like to thank Mr. John Clary, Dr. Bhim Thapa, Ms. Jade Rodgers, and Ms. Alicia Healey for excellent technical support. We are also grateful to Dr. George Hodges for providing critical review of the pathology and manuscript.

